# Subcellular mRNA localization patterns across tissues resolved with spatial transcriptomics

**DOI:** 10.1101/2025.09.07.674688

**Authors:** Roy Novoselsky, Ofra Golani, Tal Barkai, Merav Kedmi, Inna Goliand, Michal Fine, Ilan Kent, Ido Nachmany, Shalev Itzkovitz

## Abstract

Subcellular RNA localization, including nuclear retention and apical-basal compartmentalization in polarized epithelia plays a central role in post-transcriptional regulation. However, methods for high-throughput mapping of mRNA localization within intact tissue sections remain limited. Here, we apply high-resolution spatial transcriptomics (VisiumHD) to systematically resolve intracellular mRNA localization across diverse mammalian tissues. We introduce a computational approach that extracts subcellular features from spatial data and quantifies transcript localization patterns. Using this framework, we map apical-basal mRNA localization and nuclear retention in gastrointestinal epithelia and in liver hepatocytes. Our analyses reveal conserved and tissue-specific localization signatures that can be readily obtained from standard high-definition spatial transcriptomics experiments. This approach broadens the scope of spatial transcriptomics by enabling routine investigation of intracellular RNA distributions in both healthy and diseased tissues.

## Introduction

Spatial transcriptomics has revolutionized our ability to study gene expression within intact tissue environments, preserving both tissue architecture and cellular neighborhoods. However, most applications to date have focused on intercellular variation, with limited investigation of mRNA organization at the subcellular level. Understanding how transcripts are spatially distributed within cells offers unique insights into post-transcriptional regulation and functional cell polarity. This is supported by growing evidence that intracellular mRNA localization plays a critical role in mammalian cell biology (1–11). One well-characterized form of mRNA localization is apical-basal compartmentalization in epithelial cells, where specific transcripts are preferentially enriched in either the apical or basal domain(12–16). Another form of mRNA localization is nuclear retention, which is traditionally associated with splicing-defective or non-coding RNAs, but has also been observed for fully spliced transcripts in mammalian cells (17–20).

Previously, high-throughput mapping of mRNA localization within tissues has been time-consuming and technically demanding. Early approaches relied primarily on labor-intensive methods such as fractionation-based sequencing and laser capture microdissection (LCM). Recent technological advances have expanded the toolkit for studying spatial RNA localization, utilizing either targeted imaging-based techniques such as Xenium (21), MERSCOPE (22), CosMx (23) and seqFISH (24), or spatial sequencing-based spatial transcriptomics approaches, such as VisiumHD (25), Stereo-seq (26) and SlideSeq (27). Computational tools modeling data from imaging-based techniques yielded important insights into subcellular mRNA localization (28–30). However, similar tools to extract sub-cellular localization information from spatial sequencing techniques are currently lacking. Here, we develop HiVis, a user-friendly framework for spatial transcriptomics that combines advanced image-processing approaches to enable subcellular mRNA localization analyses. We systematically profile apical-basal mRNA polarization and nuclear retention across multiple mouse and human tissues using VisiumHD and demonstrate that our computational tool resolves subcellular transcript distribution and identifies polarized and nuclear-retained mRNAs in organs such as the small intestine, colon, and liver.

## Results

### Development of computational platform for VisiumHD analysis

Mapping subcellular mRNA localization across tissues requires an integrated approach that combines high-resolution image analysis with a flexible computational framework for downstream transcriptomic analysis. While tools such as *bin2cell* (31), *SpatialData* (32), Enact (33) and *Thor* (34) provide powerful frameworks at the cell or tissue scale, they do not facilitate a streamlined analysis of RNA subcellular localization.

To address this gap, we developed HiVis (HD Integrated Visium Interactive Suite), an interactive, user-friendly, end-to-end platform optimized for 10x VisiumHD datasets, but extended to support other spatial transcriptomics methods that include a registered image (Fig. S1). HiVis enables robust identification of subcellular compartments and supports application across a wide range of tissues and experimental conditions. The main features of the tool are:

- Integration with QuPath (35), providing a powerful image analysis toolkit that includes pixel classification, interactive annotation, and object feature extraction.
- Flexible, tissue-oriented segmentation and aggregation, supporting multiple levels of resolution (e.g., bins, single cells, or other objects) with algorithms such as StarDist (36), Cellpose (37), and InstanSeg (38).
- Bidirectional data transfer between bins and cells (or other objects), enabling aggregation from bin to cell level as well as redistribution of cell-level information back to bins. This allows both global cell-level analysis and fine-grained subcellular investigation.
- Comprehensive downstream analysis and visualization, with full compatibility with widely used spatial transcriptomics frameworks such as Scanpy (39) and Squidpy (40).

We applied HiVis to systematically assess mRNA localization across multiple tissues. To this end, we analyzed both published and publicly available datasets, as well as newly generated VisiumHD data from livers of two mice. Using this framework, we mapped apical-basal localization patterns in mouse small intestine, human colon, and the more complex architecture of mouse liver (Fig. 1). In addition, we profiled nuclear-cytoplasmic localization in these tissues, as well as in human liver. We validated our results using LCM and fractionated RNA sequencing (RNA-seq) datasets. To further validate our findings, we performed single-molecule fluorescence in situ hybridization (smFISH) or hybridization chain reaction (HCR-FISH).

**Figure 1.**
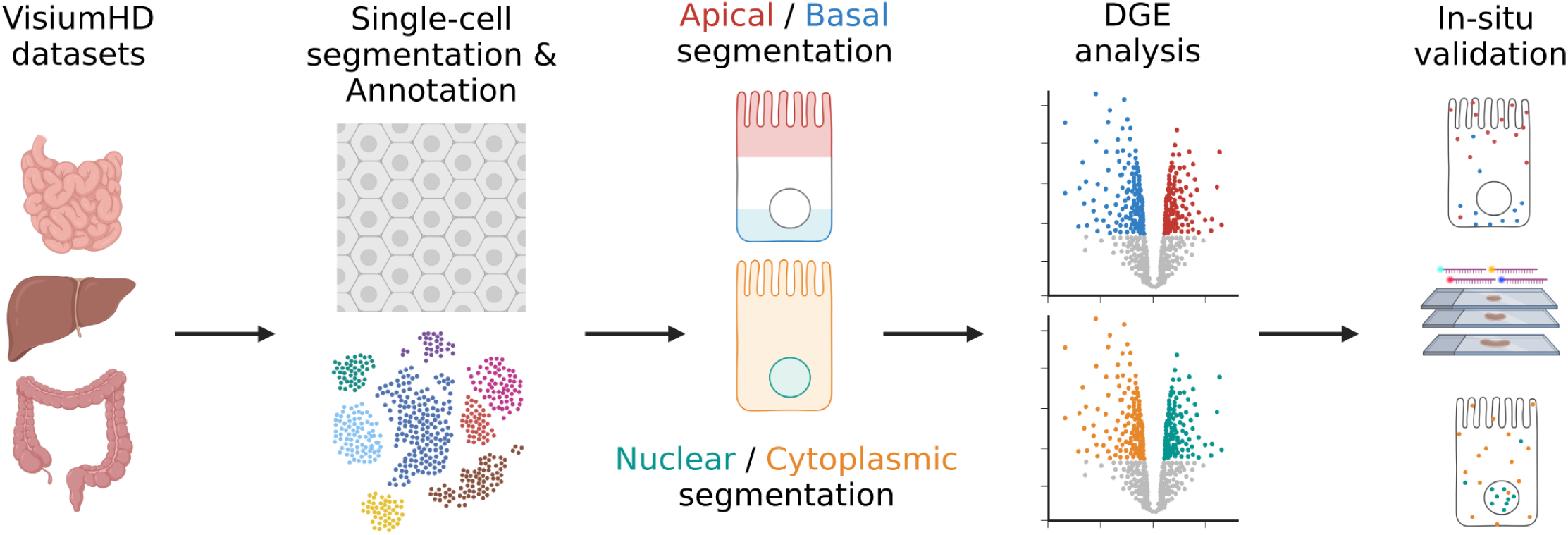
Workflow for detection of subcellular mRNA localization using HiVis. Analysis workflow for VisiumHD datasets. Images are segmented into single cells, tissue and cell type specific information is incorporated to enable precise subcellular analyses. Apical–basal and nuclear–cytoplasmic compartments are assigned to spatial bins, and differential gene expression analysis is performed to identify genes localized to these compartments. Validation is carried out using in-situ hybridization stainings. The diagram was created with BioRender.com.

### Reconstructing zonation patterns in mouse small intestine and liver using HiVis

Epithelial tissues such as the intestine and the liver exhibit pronounced zonation patterns, which strongly influence gene expression and cellular function (41–44). Delineating cell states across these spatial domains is therefore not only a critical first step in tissue-level analysis, but also an essential prerequisite for interpreting subcellular RNA localization.

To assess our ability to reconstruct zonation patterns using HiVis, we analyzed a publicly available VisiumHD dataset of the mouse small intestine (10x genomics, 2024). Because many villi were damaged in the tissue handling process (i.e. Swiss-roll preparation), and to minimize bias from proximal–distal variation (45), we restricted our analysis to the proximal small intestinal segment (Fig. S2a), which contained intact crypt-villi units. We performed single-cell segmentation (Fig. 2a) followed by aggregation of all 2×2 µm bins included in each cell, to obtain single-cell resolved data. This enabled transcriptomic analysis and cell-type assignment using standard single cell RNA-seq (scRNA-seq) workflows (Fig. 2b–d, S2b–e). The identified cell types were consistent with their expected spatial organization along the intestinal epithelium (e.g., muscle, Paneth cells, and enterocyte zones from villus base to tip).

**Figure 2.**
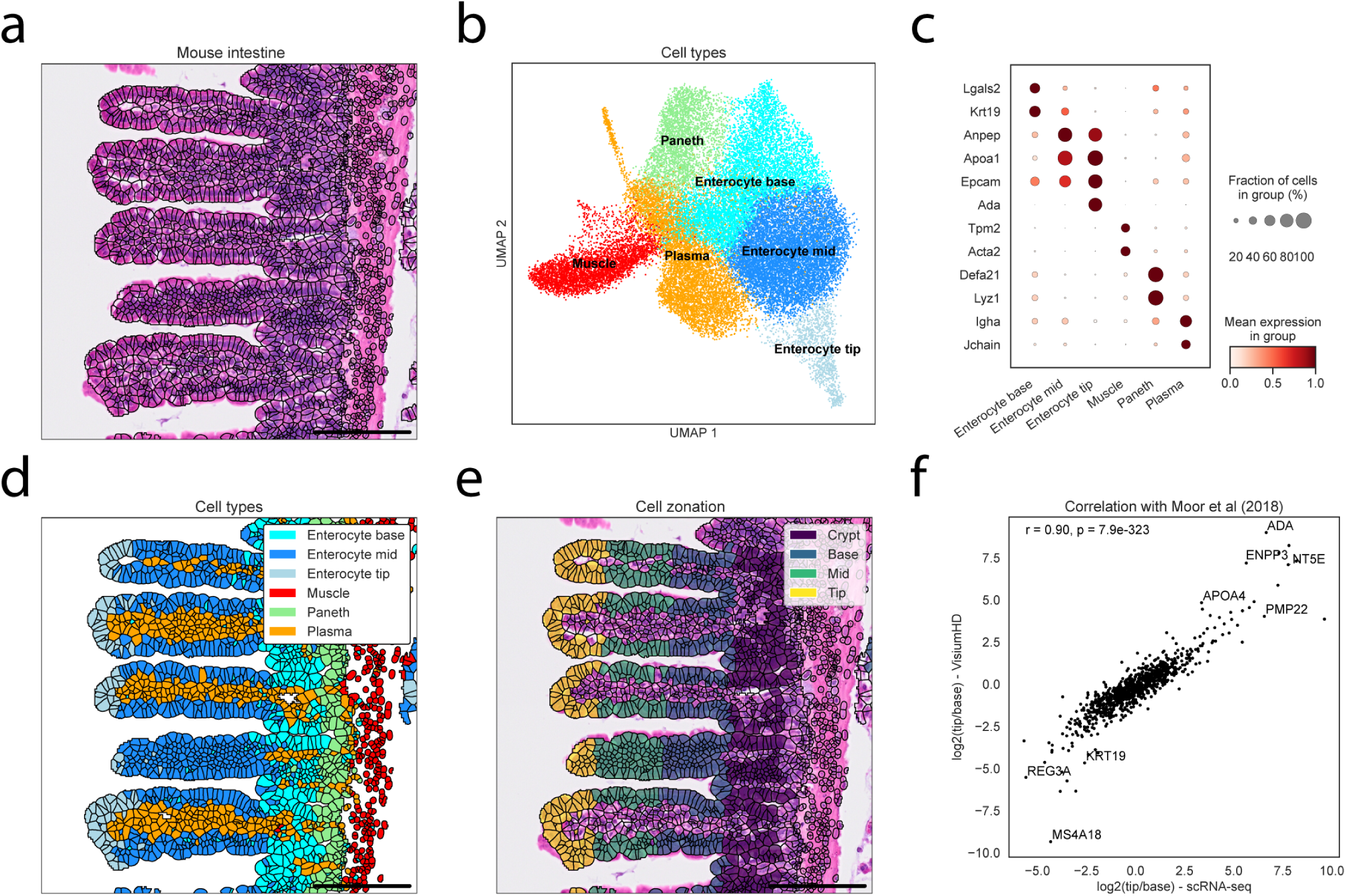
Reconstructing zonation along the crypt-villus axis in the mouse small intestine. **(a)** Single-cell segmentation of mouse small intestine VisiumHD. Scale bar 100 µm. **(b)** UMAP of single cells colored by cell type. **(c)** Dot plot of normalized gene expression for selected markers across cell types. **(d)** Same field of view (FOV) as (a), with cells colored by cell type. Colors correspond to the cluster colors in (b)**. (e)** Same FOV as (a), with cells colored by villus zones. **(f)** Spearman correlation of center of mass (COM, Methods) between the current VisiumHD analysis and previously published scRNA-seq data (41). Selected gene names are shown.

To establish spatial zonation independently of the transcriptome, we manually segmented individual villi (Fig. S2f) and identified the muscle layer using a pixel classifier (Fig. S2g). Based on this annotation, we calculated the distance of each bin and cell from the muscle layer (Fig. S2h–i). Restricting the analysis to epithelial cells, we then compared gene expression patterns between cells at the villi tips and bases (Fig. 2e, Supplementary table 1). The resulting zonation profiles showed strong concordance with previously published scRNA-seq data that was focused on zonation patterns across the villus axis (41), with a correlation of r=0.87 (Fig. 2f). To further demonstrate the utility of HiVis in extracting spatially resolved single-cell data from image-based platforms beyond VisiumHD, we applied it to a Xenium dataset of the mouse small intestine (46), revealing distinct villus zonated cellular states (Fig. S3a-f). We also demonstrated that HiVis supports other platforms by analyzing Stereo-seq dataset from mouse brain (47)(Fig. S3g-i). Our analyses confirms that HiVis can accurately reconstruct villus zonation, establishing the spatial framework required for subsequent subcellular localization analyses.

Next, we generated VisiumHD data from livers of two mice. We stained the sections with DAPI, which stains nuclei, and fluorescently-labeled antibodies against Na⁺/K⁺-ATPase (ATP1A1) and CD31 to facilitate identification of nuclear, membrane and basal (sinusoidal-facing) cellular compartments respectively (Fig. 3a-b, S4a). Using HiVis, we segmented hepatocytes (parenchymal cells) and non-parenchymal cells (NPC) independently, based on regions depicted by a pixel classifier (Fig. 3b, S4b, Methods). We then aggregated the included 2×2 µm bins to obtain single cell transcriptomics, and reconstructed hepatocytes zonation maps based on established landmark genes (Fig. 3c, S4c, Methods), as previously described (48). From this, we derived zonation profiles for each gene, which were highly consistent across the two samples (Fig. S4d, Supplementary table 1). These profiles showed strong positive correlations with previously published zonation profiles that were based on either spatially-resolved scRNA-seq (43) or high-definition spatial transcriptomics data obtained using Stereo-seq (49) (Fig. 3d).

**Figure 3.**
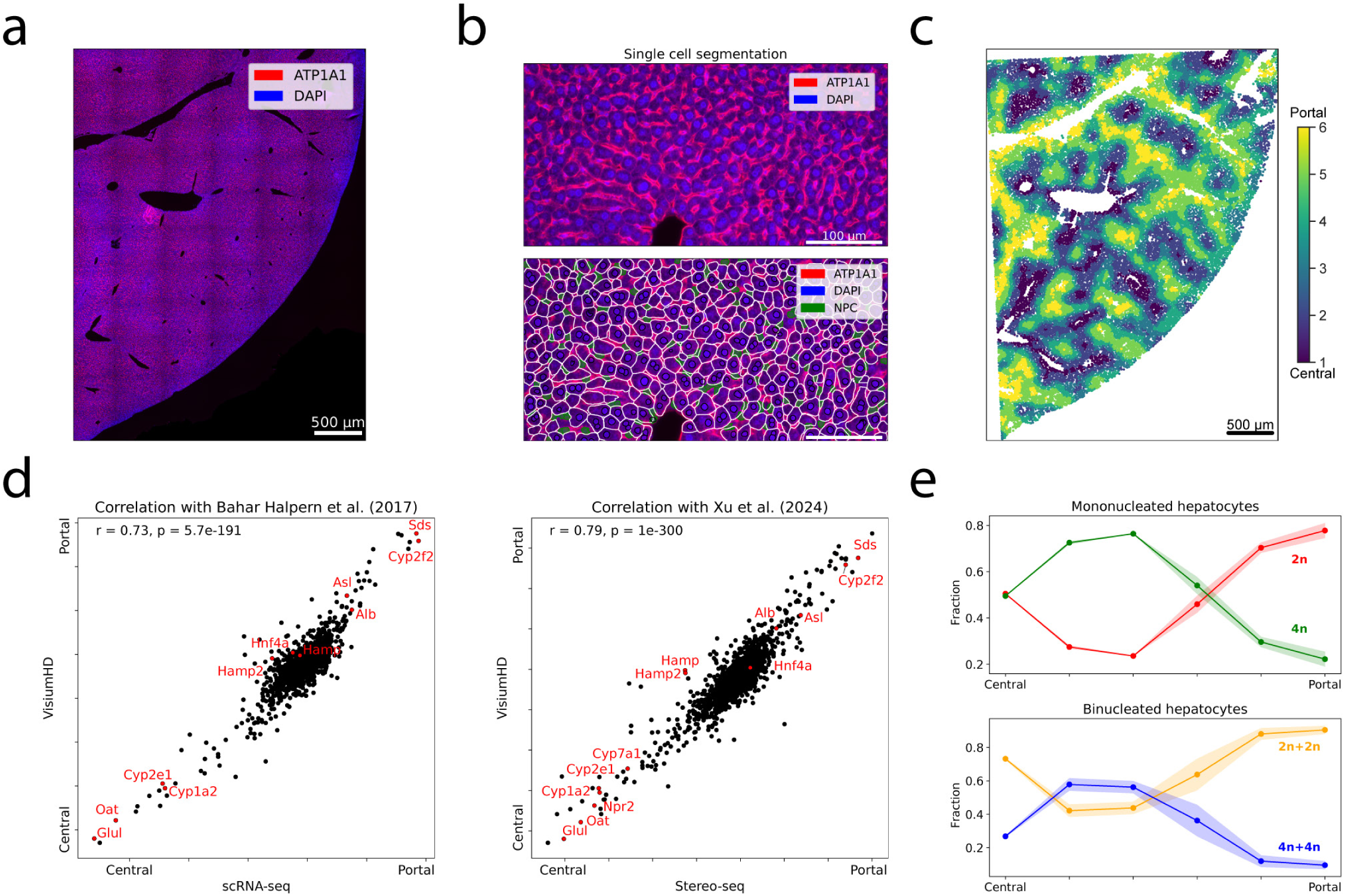
Reconstructing hepatocyte zonation across liver lobules. **(a)** VisiumHD immunofluorescence image of mouse liver. Representative image of 2 samples. **(b)** Mouse liver VisiumHD immunofluorescence image (top) and overlayed with single-cell segmentation (bottom). NPC – non parenchymal cells, residing in the sinusoids, which are segmented based on CD31 and ATP1A1 staining (Methods). **(c)** Single-cell VisiumHD data of mouse liver, colored by zones along the central-portal axis. Shown is a representative plot from one mouse. **(d)** Spearman correlation plots of center of mass across liver zones based on current VisiumHD and previously published scRNA-seq (52) (left) and Stereo-seq spatial transcriptomics (49) (right). **(e)** Mean fraction of ploidy classes in both mononucleated (top) and binucleated (bottom) hepatocytes across the liver lobule in both mice. Patch is standard error of the mean.

As our single-cell segmentation was based on membrane-staining, we could use morphological features extracted from the single-cell segmentation. We found that as previously shown (50), the overall sizes of hepatocytes decreased significantly from the central vein towards the portal vein (r=-0.26, p<1e-300, Fig. S4e). Hepatocytes in mice contain either one or two nuclei, with distinct ploidy levels that scale with nuclear size (51). Using the number of nuclei and their sizes, we assigned a ploidy class for each cell (Fig. S4f). In concordance with previously published results (51) we found that hepatocytes in the mid-lobule zones in mice exhibited higher ploidy compared to central and portal hepatocytes (Fig. 3e). These results demonstrate that HiVis can accurately reconstruct lobular zonation in the liver. We next turned to investigate subcellular RNA localization in zonally stratified cells in these two stereotypically zonated tissues.

### Polarized mRNA distribution in the intestinal epithelium

Following the cell-level analysis of epithelial cells in the mouse intestine, we next investigated the subcellular distributions of mRNAs. The intestinal epithelium is composed of an apical side facing the nutrient and microbial enriched lumen, and a basal side facing the blood stream. These sides differ in morphology (53,54), protein (13,55) and even RNA content (13,14,16). To this end, we assigned apical and basal identities to 2×2 µm bins using a pixel classifier (Fig. 4a). Differential gene expression (DGE) analysis revealed transcripts significantly enriched in either the apical or basal domains (Fig. 4b, Supplementary table 2). The apical-basal enrichment patterns obtained with pixel classifier were consistent with those derived from manual annotations (r=0.83, p=6e-85, Fig S5a). We validated the apical polarization of Apob and the basal polarization of Net1 using smFISH (Fig. 4c). The resulting polarization profiles showed strong concordance with previously published data obtained by LCM of apical and basal compartments (14) (Fig. 4d). We next examined whether apical-basal polarization varies along the villus axis and found that profiles at the tip and base were largely similar, consistent with previous observations (14) (Fig. S5b). We further examined whether apically localized transcripts exhibit preferential luminal versus perinuclear enrichment (Fig. S5c–d) and observed clear differences between these patterns (Supplementary Table 2). For example, *Lct* showed strong luminal apical localization, whereas *Atp1b1* displayed a more uniformly distributed apical pattern (Fig. S5e–g).

**Figure 4.**
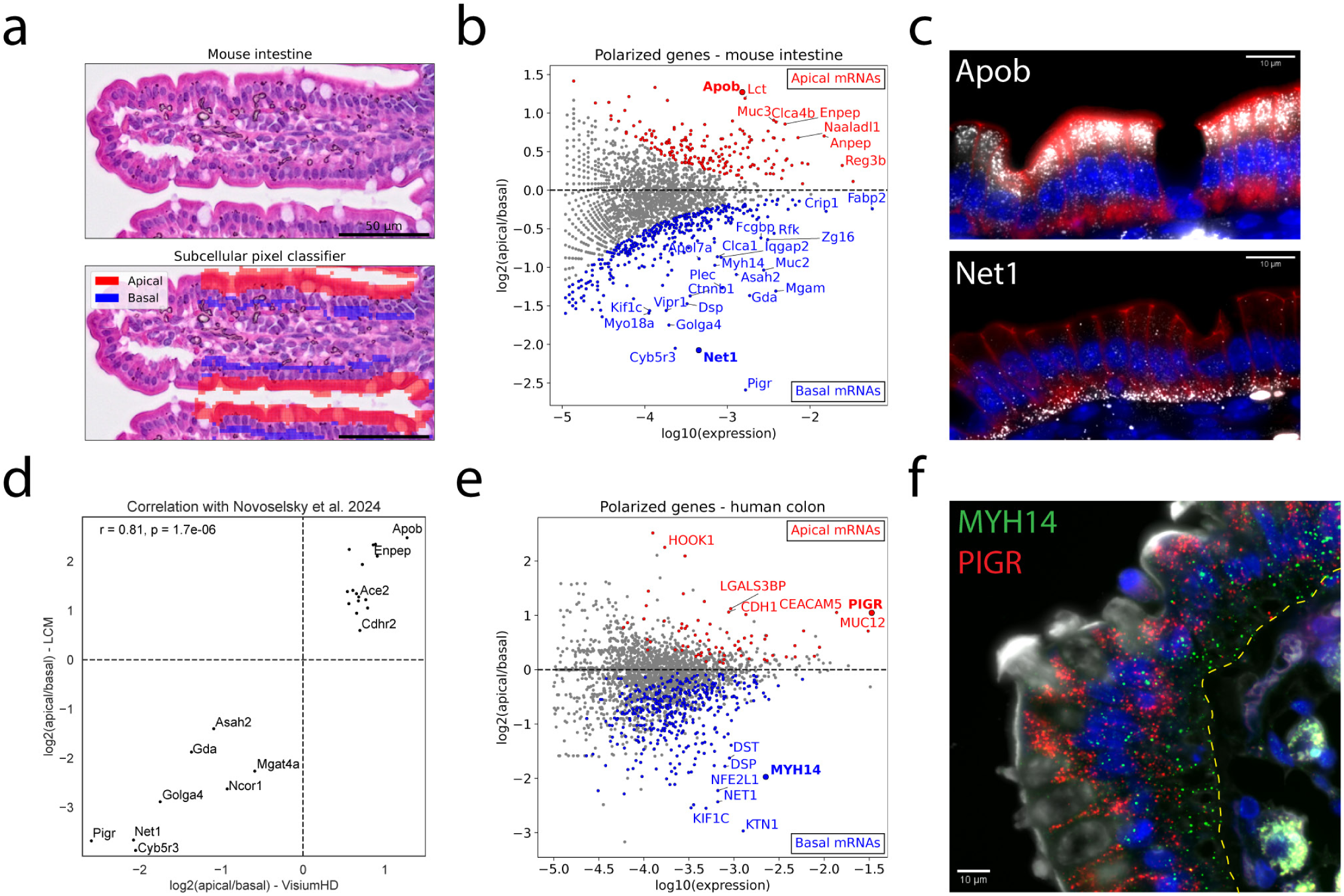
Mapping apical-basal mRNA localization in mouse small intestine and human colon. **(a)** Mouse small intestine VisiumHD H&E image (top) and overlayed with 2×2µm bins colored by apical/basal assignment, based on a pixel classifier (bottom). **(b)** MA-plot of differential gene expression results in mouse proximal small intestine. Significant genes (q<0.05) are colored red and blue, bold genes shown in smFISH validation in (c). Selected gene names are shown. **(c)** smFISH staining of Apob (top) and Net1 (bottom) in mouse proximal small intestine. Red staining is E-cadherin which stains epithelial basolateral membranes, blue staining is DAPI. Scale bar, 10 µm. **(d)** Spearman correlation of apical/basal bias based on current VisiumHD and previously published scRNA-seq (14). Only genes with q<0.25 in both datasets are shown. **(e)** MA-plot of differential gene expression in human colon. Significant genes (q<0.25) are colored red and blue, bold genes shown in HCR-FISH validation in (f). Selected gene names are shown. **(f)** HCR-FISH staining of PIGR and MYH14 in human colon. White staining is Wheat Germ Agglutinin (WGA), blue staining is DAPI. Dashed yellow line marks the basal side of the epithelial cells, depicted by WGA staining. Images in (c) and (f) are representative of 2 individuals. Scale bar, 10 µm.

These results demonstrated that our approach can accurately capture apical-basal polarization in intestinal epithelial cells, prompting us to extend the analysis to human colon tissue, a tissue in which epithelial mRNA polarization has not been systematically explored. To this end, we analyzed published VisiumHD data from five patients with colorectal cancer (25), focusing on adjacent normal or control regions. As in the small intestine, we classified bins into apical and basal epithelial compartments and performed DGE analysis (Fig. S5h) and identified transcripts enriched in each domain (Fig. 4e, Supplementary table 2). Correlation analysis of polarization profiles between colon and previously published jejunum data revealed highly similar patterns across the two organs (Fig. S5i). Apical transcripts included *PIGR*, *CDH1*, and *CES2*, whereas basal transcripts included *KTN1*, *NET1*, and *MYH14*. We validated *MYH14* and *PIGR* by HCR-FISH (Fig. 4f). As shown in previous work (14), PIGR exhibited significant apical localization in humans, and significant basal localization in mice (Fig. 4b, d, e, f). Together, these results demonstrate that colon epithelial cells exhibit clear apical–basal mRNA polarization, extending this localization pattern to an additional segment of the gastrointestinal tract.

### Specific mRNAs are basally localized in mouse hepatocytes

Having demonstrated that HiVis can resolve apical-basal transcript localization in the simple columnar epithelium of the digestive tract, we next turned to a more complex tissue: the mouse liver. Hepatocytes in the mammalian liver are arranged in radial sheets that span the portal-central axis. Each hepatocyte has two basal sides, facing the sinusoidal blood vessels, and cortical sides that face adjacent hepatocytes (these include the apical sides that face the bile canaliculi) (56). In this case, apical and basal domains are not easily identifiable and therefore cannot be reliably isolated by LCM. As a result, mRNA localization to these compartments has remained uncharacterized, making it an ideal test case for our approach. Our VisiumHD data included staining for ATP1A1 and CD31, which marked membranes and sinusoidal endothelial cells respectively. Using these markers, we identified sinusoids between hepatocytes (Fig. 5a) and calculated the distance of each bin within hepatocytes to both the nearest sinusoid and to the cell border (Fig. 5b). This enabled assignment of cortical (close to any membrane regardless of distance from sinusoids) and basal (close to sinusoids) identities to each bin (Fig. 5c–d). Differential gene expression analysis identified several basally enriched transcripts, including *Scd1*, *Pigr*, *Cald1*, and *Cyb5r3* (Fig. 5e, Supplementary table 2). Validation by smFISH confirmed basal localization of *Pigr* and *Cyb5r3* (Fig. 5f), providing, to our knowledge, the first evidence of cytoplasmic mRNA polarization in hepatocytes. These findings demonstrate that HiVis can be applied to VisiumHD data from complex tissues to resolve subcellular mRNA localization.

**Figure 5.**
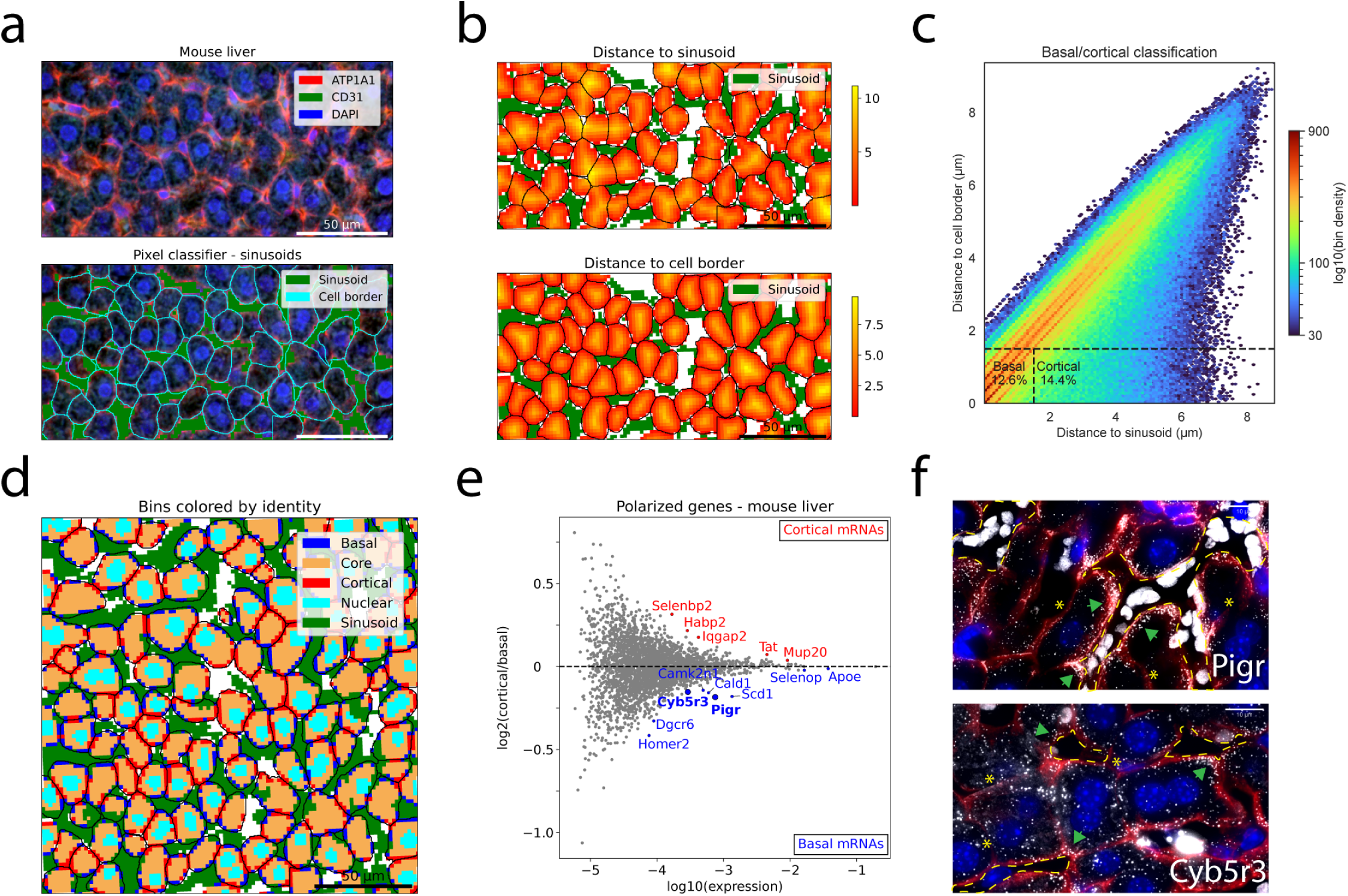
Mapping basal mRNA localization in mouse liver. **(a)** Mouse liver VisiumHD immunofluorescence image (top) and overlayed with 2×2µm sinusoid bins, based on a pixel classifier and with cells segmentation (bottom). **(b)** Same FOV as (a), with bins colored by distance to sinusoids (top) and distance to cell border (bottom). Distance is in µm. **(c)** Representative density plot of 2×2µm bins from one mouse liver. Black squares indicate the bins that were classified as cortical or basal. **(d)** 2×2µm bins in mouse liver, colored by intracellular location. **(e)** MA-plot of differential gene expression in mouse liver. Significant genes (q<0.25) are colored red, blue genes are shown in smFISH validation in (f). Selected gene names are shown. **(f)** smFISH staining of Pigr (top) and Cyb5r3 (bottom) in mouse liver. Red staining is ATP1A1, blue staining is DAPI. Dashed yellow line – outlines sinusoids between hepatocytes. Green arrows point to the basal sides of hepatocytes, yellow asterixis depict cortical sides. Image representative of 2 animals. Scale bar, 10 µm.

### Mapping nuclear and cytoplasmic mRNA distribution in multiple tissues

We next used our HiVis computational pipeline and spatial transcriptomics datasets to extract the relative distribution of mRNAs between the nucleus and cytoplasm. As a starting point, we analyzed the mouse liver (Fig. 6a), where nuclear retention of transcripts has been previously described using cell fractionation followed by RNA-seq (18). We identified transcripts enriched in either the cytoplasmic and nuclear compartments (Fig. 6b, Supplementary table 2). Cytoplasmic genes included the basally localized transcripts identified earlier, *Pigr* and *Scd1*, along with key hepatocyte markers such as *Apoc1*, *Apoa2*, *Apoe*, and *Alb*. Nuclear-enriched transcripts included *Mlxipl*, *Nlrp6* and *Sema4g*. Correlation analysis revealed a moderate positive agreement with the results of Bahar Halpern et al. (18) (r=0.66, p=3e-103, Fig. 6c), despite major differences in methodology, including bulk versus spatial resolution, poly(A) versus targeted capture, and overall sequencing strategies. Utilizing the ability to analyze zonal hepatocytes, we asked whether nuclear localization patterns vary across the liver lobule radial axis. We did not observe major differences in nuclear-cytoplasmic bias between central and portal regions, indicating that hepatocyte nuclear retention is consistent across the lobule (Fig. S6a). These findings validate that HiVis can reliably resolve nuclear-cytoplasmic mRNA distribution from VisiumHD data.

**Figure 6.**
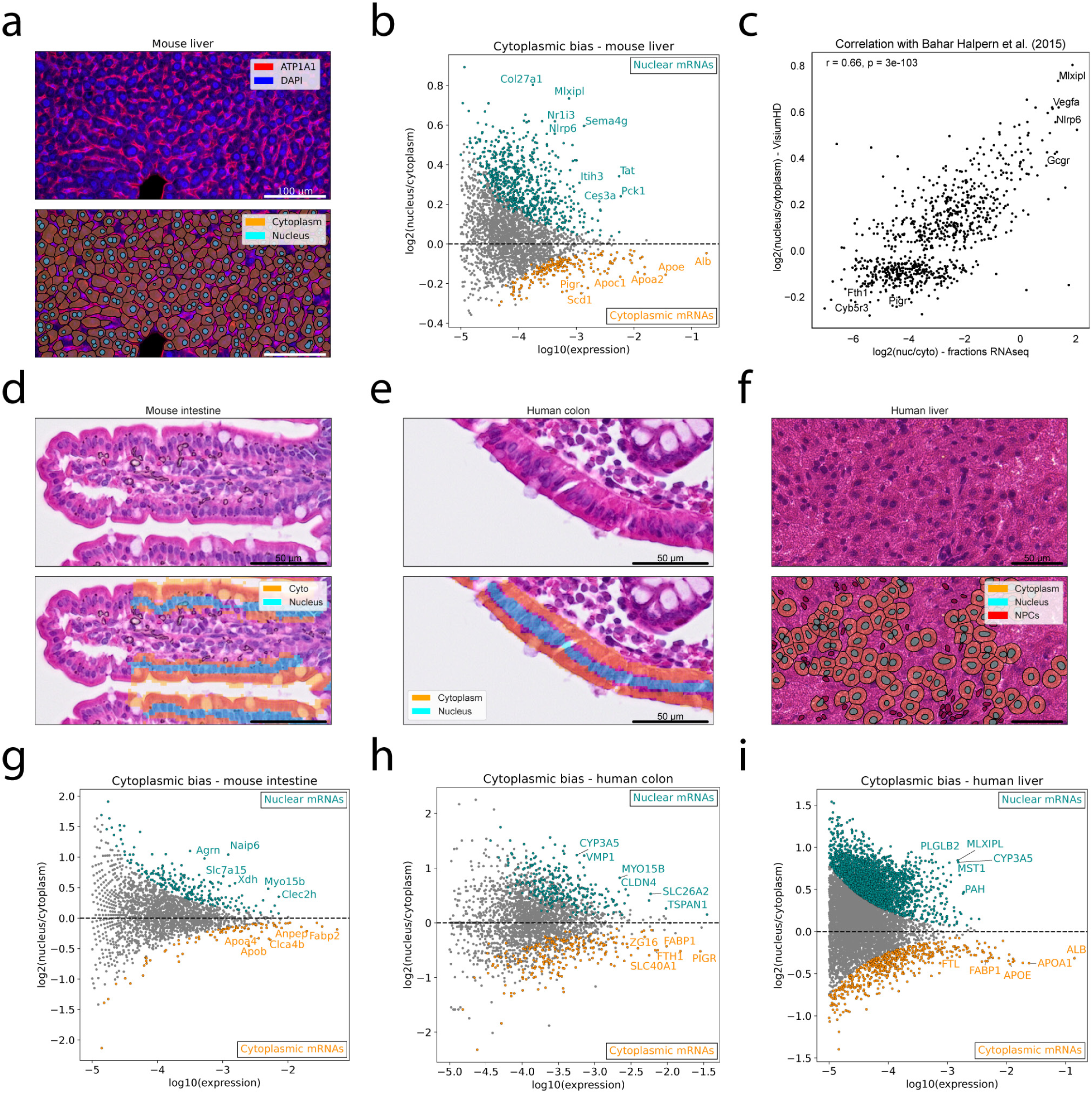
Mapping nuclear retention across tissues. **(a)** Mouse liver VisiumHD immunofluorescence image (top) and overlayed with hepatocyte cellular and nuclear segmentation (bottom). **(b)** MA-plot of differential gene expression results in mouse liver. Significant genes (q<0.01) are colored red. Selected gene names are shown. **(c)** Spearman correlation plots of cytoplasmic/nuclear bias based on current VisiumHD and previously published RNA-seq (18). **(d, e, f)** Top – H&E images from VisiumHD of mouse proximal small intestine (d), human colon (e) and human liver (f). Bottom – same images as top, overlayed with 2×2µm bins colored by nuclear/cytoplasmic assignment. **(g, h, i)** MA-plot of differential gene expression results in mouse intestine (g), human colon (h) and human liver (i). Significant genes are colored red (g: q<0.05, h: q<0.25, i: q<0.05). Selected gene names are shown.

High-throughput mapping of nuclear–cytoplasmic mRNA localization has previously been explored in the mouse liver, yet not in the gastrointestinal tract nor in human. Therefore, we extended our analysis to additional tissues, including mouse proximal small intestine (Fig. 6d), human colon (Fig. 6e), as well as a recently published human liver dataset from a live organ donor (44) (Fig. 6f, S6b). While performed only in mouse liver, comparable analyses have not been carried out in these tissues. We used the H&E image to automatically extract the nuclear and cytoplasmic compartments. In each tissue, we detected transcripts preferentially localized to either the nuclear or cytoplasmic compartments (Fig. 6g-i, Supplementary table 2). We found that nuclear-cytoplasmic bias was conserved between mouse and human (Fig. S6c, r=0.71, p=1.7e-31), as well as between human liver and colon (Fig. S6d, r=0.74, p=1.3e-20). We applied similar analysis on published Xenium dataset of mouse small intestine, to further demonstrate the compatibility of HiVis with other platforms. We confirmed that nuclear genes such as Neat1 and Vegfa were enriched in the nucleus, while Apoa1 and Fabp2 were enriched in the cytoplasm (Fig. S6e-f). These findings establish HiVis as a versatile framework for characterizing nuclear-cytoplasmic mRNA localization in diverse tissue contexts.

## Discussion

mRNA localization is a widespread but incompletely understood phenomenon, particularly in epithelial cells (11). Although specific cases have been shown to be important for normal cell function (16), their extent across cell types and tissues remain unclear. With advances in high-resolution spatial transcriptomics, we set out to systematically map mRNA localization across tissues using the 10x VisiumHD platform. Applying this approach, we successfully characterized both apical-basal and nuclear-cytoplasmic biases in several tissues of the digestive system from mouse and human (Fig. 1). To enable these analyses, we developed HiVis, a flexible tool that streamlines analysis across diverse datasets by integrating image processing tools, tissue-specific single-cell segmentation, customized workflows, and subcellular transcript-localization analyses. In contrast to existing frameworks for subcellular transcriptomics, such as SPRAWL (28), Bento (29) and Ella (30), HiVis does not rely on isolated single-cell masks as the primary spatial unit. Instead, it integrates image-derived tissue context with segmentation outputs, allowing polarity features that depend on anatomical structure – such as lumen proximity in the intestine or sinusoidal orientation in the liver – to be captured. HiVis can also analyze binned spatial data when segmentation is incomplete, while still supporting full mask-based workflows when high-quality masks are available. Together, these features provide a unified framework for linking anatomical annotation, spatial zonation, and subcellular localization.

Our results showed strong concordance with previously published datasets, including LCM-derived profiles of apical and basal compartments from mouse intestinal epithelium and fractionated RNA-seq of nuclear and cytoplasmic compartments from mouse liver. Together, these comparisons support the accuracy of our approach for mapping global mRNA localization. Unlike the intestinal epithelium, the distinct morphology of hepatic plates complicates the characterization of apical–basal mRNA polarization in the liver. In this study, we combined membrane and endothelial staining with high-resolution spatial transcriptomics to identify, for the first time, a small set of hepatocyte transcripts exhibiting basal polarization. Our analyses further enabled comparison of mRNA localization across species and organs. Most localization patterns appeared conserved in both contexts. Moreover, although cells along tissue spatial axes – such as villi in the small intestine and lobules in the liver – show pronounced differences in expression levels, we observed little variation in mRNA localization across these zones. These findings suggest that under normal conditions, the subcellular localization of most mRNA species is largely preserved in epithelial cells of the digestive system.

High-definition spatial transcriptomics has advanced rapidly in recent years, enabling increasingly detailed studies of gene expression within intact tissues. Although imaging-based approaches provide higher spatial resolution, they are often challenging to implement due to large data volumes, complex computational requirements, and most importantly – the limited number of targeted genes, which introduces bias. Sequencing-based spatial transcriptomics, in contrast, is highly scalable and continues to improve, though it remains constrained by molecular diffusion and spatial resolution. For example, the 2 µm resolution of VisiumHD can blur adjacent subcellular regions, causing perinuclear cytoplasmic transcripts to be misclassified as nuclear. Its major advantages include unbiased whole-transcriptome coverage and compatibility with FFPE samples, enabling exploratory analyses of archival and diseased tissues under diverse conditions. Building on these strengths, our approach adds a new layer of analysis that can reveal mRNA localization across tissues, cell types, and physiological or pathological contexts.

## Methods

### Experimental methods

#### Animal experiments and tissue preparations

Mouse experiments were approved by the Institutional Animal Care and Use Committee of the Weizmann Institute of Science (approval number 05220824–2) and performed in accordance with institutional guidelines. Two-months-old C57BL/6 male mice, housed under regular 12h light-dark cycle and fed ad libitum were used in the experiments. Mice were sacrificed by cervical dislocation and tissues were collected and fixed in 4% PFA in PBS in 4°C. Livers were incubated for 24 h, transferred to 70% Ethanol, embedded in paraffin, and stored at room temperature. Proximal jejunums were incubated for 3h, followed by overnight fixation in 4% FA + 30% sucrose at 4°C under gentle rotation. The jejunums were rolled to a “Swiss-rolls”, embedded in OCT, and stored at −80°C.

#### Mouse liver VisiumHD experiment

VisiumHD spatial transcriptomics was performed on liver tissue sections from two male C57BL/6 mice (2.5 months old), following the official 10x Genomics protocol.

##### Immuno staining and fluorescence microscopy

Formalin-fixed, paraffin-embedded (FFPE) tissue blocks were sectioned into 5-μm slices and mounted onto poly-L-lysine-coated glass slides (Sigma-Aldrich, P0425).Tissue sections were immunostained according to the VisiumHD user guide. Primary antibodies included mouse anti-CD31 (ab28364, 1:100), rabbit anti-ATP1A1 (ab76020, 1:1000), and rabbit anti-LMNA (ab26300, 1:100), incubated for 90 minutes. Secondary antibodies were applied for 20 min, including donkey anti-mouse Cy3 (Jackson ImmunoResearch, 715-066-151, 1:200), donkey anti-rabbit Cy5 (Jackson ImmunoResearch, 711-175-152, 1:200), and DAPI (100 ng/mL, Sigma-Aldrich, D9542). Tissue sections were mounted in 85% glycerol in RNase-free water and covered with No. 1.5 coverslips (Marienfeld, 0102222). Fluorescence imaging was carried out on a widefield Leica DMi8 inverted microscope (Leica Microsystems Heidelberg GmbH, Germany) operated with LAS X software. The system was equipped with a Leica 12-bit monochrome camera (DFC7000GT-0127092118) and a multi-band fluorescence filter cube (QWF-S-T). Images were acquired with an HC PL APO 20×/0.80 dry objective (11506530) at a pixel size of 0.323 µm. A Tile Scan (XY mode) was performed using the LAS X Tile Scan function, and the resulting tiles were stitched in LAS X.

##### Library preparation and sequencing

Targeted transcript capture and amplification were carried out using the Mouse Probe Set v2 (10x Genomics). Ligated probes were transferred to VisiumHD spatial barcoding slides using CytAssist (v2.1.0.14) at 37 °C for 30 minutes, following the manufacturer’s instructions. Library preparation was conducted using the VisiumHD Spatial Gene Expression User Guide, including 8 cycles of PCR for cDNA amplification. Prepared libraries from each slide were loaded onto individual lanes of an Illumina NovaSeq X flow cell at a concentration of 150 pM and sequenced.

#### Colon tissue collection and ethics declaration

Colon samples were obtained from patients undergoing colectomy for malignancy. All surgeries and FFPE block preparation were performed at Sheba Medical Center, Israel. This study was approved by the Sheba Medical Center Helsinki committee (approval number SMC-8665-21). All participants provided written informed consent prior to inclusion. The study was conducted in accordance with the Declaration of Helsinki and relevant local regulations

#### Single molecule fluorescence in situ hybridization

Mice Jejunums and livers were stained as previously described (57). Probe libraries were designed using the Stellaris FISH Probe Designer Software (Biosearch Technologies, Inc., Petaluma, California, United States of America). Probe libraries were coupled to Cy5, or Alexa 594. Cryosections (6 μm) of fixed mouse Jejunum or liver were mounted on poly-L-lysine pre-coated coverslips and used for probe hybridization. Sections were fixed for 15 min in 4% FA in PBS, washed with PBS, and incubated for 2 h in 70% ethanol in 4°C. Sections were washed with 2× SSC (Ambion AM9765) for 5 min, then permeabilized for 10 min with proteinase K (10 µg/ml, Ambion AM2546) followed by 2 washes with 2× SSC (Ambion AM9765) for 5 min. Tissues were incubated in wash buffer (Jejunum with 20%, livers with 15% Formamide Ambion AM9342, in 2× SSC) for 10 min in a 30°C incubator. Tissues were incubated with hybridization mix mixed with probes (Jejunum with 20%, livers with 15% Formamide, 10% Dextran sulfate Sigma D8906, 1 mg/ml *E*. *coli* tRNA Sigma R1753, 10% 20× SSC, 0.02% BSA Ambion AM2616, 2 mM Vanadylribonucleoside complex NEB S1402S, in Rnase-free water). For Jejunum sections – FITC-conjugated anti-CDH1 antibody (1:100, BD Biosciences cat. 612131) was added to hybridization mix, and for liver sections – anti-ATP1A1 antibody (1:1500, Abcam ab76020). Samples were incubated overnight in 30°C. After the hybridization, tissues were washed with wash buffer for 30 min in 30°C, and with wash buffer with DAPI (50 ng/ml, Sigma-Aldrich, D9542) for 10 min. Liver samples were incubated with Cy3 donkey anti-rabbit secondary antibody (1:200, Jackson Immuno Research, 711-165-152) together with DAPI for 20 min. Finally, tissues were washed twice with 2× SSC, and coverslips were mounted with ProLong Gold antifade reagent (Invitrogen P36934). Images were acquired using 100× magnification. All images were performed on a Nikon-Ti-E inverted fluorescence microscope using the NIS element software AR 5.02.03.

#### HCR-FISH

Colon samples were stained according to the Molecular Instruments HCR GOLD protocol for FFPE tissues. In brief, FFPE sections were deparaffinized and rehydrated through a graded ethanol series (Xylene, Xylene, 100% Ethanol, 100% Ethanol, 95% Ethanol, 70% Ethanol, 50% Ethanol, PBS). Sections were permeabilized for 10 min with 0.5% TritonX-100 (Sigma Aldrich) followed by two washes with PBS. Tissues were incubated in hybridization buffer (HiFi HCR hybridization buffer with 100 μg/ml salmon sperm DNA, Sigma-Aldrich D7656) for 10 min in a 37 °C incubator. Tissues were incubated with hybridization buffer mixed with probes (1:50) for 16 h in 37 °C. After the hybridization, tissues were washed with HiFi HCR wash 15 min in 37 °C for 4 times. The samples were incubated with amplification buffer (HCR amplification buffer with 100 μg/ml salmon sperm DNA) for 30 min at room temperature. HCR-hairpins (H1 and H2 hairpins of X1-546, X3-647) were heated for 90 s at 95 °C and cooled in dark for 30 min. The samples were incubated with AB with 6 pmol of each hairpin for 60 min in the dark, and then rinsed twice in GOLD HCR wash buffer for 15 min. The samples were incubated with GOLD HCR wash buffer with DAPI (50 ng ml^−1^, Sigma-Aldrich, D9542) for 5 min. Additional wash with GOLD HCR wash buffer was performed, and slides were mounted with Immu-Mount (Epredia, 990402). Images were acquired using 90× magnification.

All images were performed on a Nikon-Ti-E inverted fluorescence microscope using the NIS element software AR 5.02.03.

### Computational methods

All analysis was performed using the HiVis tool (<u>V0.9</u>) in Python 3.10 and QuPath <u>0.6.0</u> (see <u>code</u> <u>availability</u>). Pixel classifiers in Qupath were trained using user-annotated examples for each class and relied solely on QuPath’s built-in multiscale image features; No manually defined landmarks or distance-based features were used.

#### Mouse intestine (VisiumHD)

Publicly available VisiumHD mouse intestine dataset (C57BL/6 mouse, Cell Ranger v3.0, 10x Genomics, 2024, March 25) was imported, analyzed, and visualized in HiVis.

Single-cell segmentation and pixel classifiers

Single-cell segmentation was conducted in QuPath (35) using the Stardist extension (36) with the “he_heavy_augment” pretrained model and the following parameters:

- Threshold: 0.5
- Normalize Percentiles: 1, 99
- Cell expansion: 2 µm in muscle regions (defined by a pixel classifier) and 7 µm elsewhere.

Before nuclear expansion, areas without tissue (defined by a pixel classifier) were excluded, and nuclei identified as artefacts by an object classifier were removed. Pixel classifiers were applied with the following settings:

- Classifier: Random trees
- Resolution: Moderate (2.19 µm/pixel)
- Features:
  - Channels: Hematoxylin, eosin
  - Scales: 1,2,4
  - All available features

##### Cell type annotations

Standard scRNA-seq analysis was performed with Scanpy (39). Genes expressed in fewer than 3 cells were removed. Data were normalized and log-transformed, and the 3000 most highly variable genes were identified and retained. Gene expression values were scaled, and PCA was performed. A neighbors graph (n_neighbors=15) and Uniform Manifold Approximation and Projection (UMAP) (58) were computed on the first 30 PCs, and Leiden clustering was applied with a resolution of 0.5. Cell types were assigned to clusters based on known marker genes (Fig. S2c).

##### Villus zonation analysis

Epithelial-specific genes were defined using a signature matrix incorporating all major intestinal cell types, primarily based on Bahar et al. (59).Fibroblast markers were taken from Bahar et al. (60), and endothelial markers from Kalucka et al. (61):

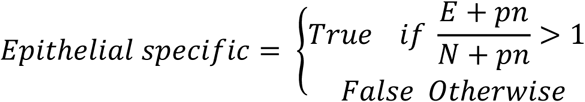

Where *E* is the maximum expression in the Enterocytes and goblet cells, *N* is the maximum expression across all non-epithelial cell types, and *pn* the smallest non-zero value in *E*.

To quantify zonation along the villus axis, a pixel classifier was used to distinguish muscle from the remaining tissue. Individual villi were then manually segmented in QuPath. For each villus, the median coordinates of the muscle-associated bins were determined, and the distance of each bin from this reference point was calculated. Zonation bins were manually defined based on their distance from muscle:

- Base villus: 400–650 µm
- Mid villus: 650–900 µm
- Tip villus: ≥900 µm

Proximal small intestine, which had full undamaged villi was segmented manually in QuPath.

For correlation with previous tip/base scRNA-seq experiment (41), the published data was imported and tip and base were assigned as zones 1-2 and 5-6 respectively. In the VisiumHD data, tip and base identities were defined in cells from the epithelial cluster, in the <u>proximal</u> region. Spearman correlation was calculated on epithelial-specific genes with maximum normalized expression > 10^-4^ in both datasets.

##### Identification of basal, apical and nuclear genes

Apical, basal and nuclear identities were assigned to bins using a pixel classifier trained with the same parameters as above, except including scale 0.5. Apical and basal genes were also classified as cytoplasmic. Differential gene expressions between apical and basal, luminal-apical and perinuclear-apical, or nuclear and cytoplasmic bins was tested using Fisher’s exact test. To avoid zonation bias, the analysis was restricted to the proximal intestine, limited to the base and mid villus zones, and focused on epithelial-specific genes. For correlation with previous apical/basal LCM experiment (14), data were imported, and Spearman correlation was calculated on genes with q-value < 0.25, mean normalized expression > 10^-5^ and |log_2_(*FC*)| ≥ 0.5. For correlation of apical/basal bias between the tip and base of villi, the same test was applied to bins in these regions, and Spearman correlation was calculated on genes with q-value < 0.05 and mean normalized expression > 10^-5^.

##### Quantification of smFISH intensity of apical genes

Luminal-apical and perinuclear-apical regions were manually segmented from epithelial cells located in spatially distant villi from two mice, based on nuclear and membrane staining, using ImageJ (62). Mean smFISH intensities for *Lct* and *Atp1b1* were quantified within each compartment, and background signal was subtracted using measurements from adjacent non-cellular regions. The ratio of luminal-to-perinuclear background-subtracted intensities was then calculated for each cell, and statistical significance was assessed using a Wilcoxon signed-rank test.

#### Mouse intestine (Xenium)

Publicly available Xenium mouse intestine dataset (46) was imported, transcripts were aggregated into 3µm bins and analyzed and visualized in HiVis.

Single-cell segmentation and pixel classifiers

Single-cell segmentation was conducted in QuPath (35) using the Cellpose extension (37,63) with the “cyto3” pretrained model with diameter set to 15µm.

Lumen annotations were segmented with a pixel classifier with the following settings:

- Classifier: Random trees
- Resolution: Moderate (1.7 µm/pixel)
- Features:
  - Scales: 1,2,4,8
  - All available features

##### Cell type annotations

Standard scRNA-seq analysis was performed with Scanpy (39). Data were normalized and log-transformed and scaled, and PCA was performed. A neighbors graph (n_neighbors=15) and UMAP were computed on the first 20 PCs, and Leiden clustering was applied with a resolution of 0.5. Cell types were assigned to clusters based on known marker genes (Fig. S3f).

##### Identification of nuclear and cytoplasmic genes

Differential gene expressions between nuclear and cytoplasmic bins were tested using Fisher’s exact test.

#### Mouse brain (Stereo-seq)

Previously published Stereo-seq mouse brain dataset (47) was imported, transcripts were aggregated into 5µm bins and analyzed and visualized in HiVis. Cells were segmented using Qupath (35) with default parameters and 2µm expansion. Standard scRNA-seq analysis was performed with Scanpy (39). Data were normalized and log-transformed and scaled, and PCA was performed. A neighbors graph (n_neighbors=15) and UMAP were computed on the first 5 PCs, and Leiden clustering was applied with a resolution of 0.8. Cell types were assigned to clusters based on known marker genes

#### Human colon

Published VisiumHD data of human colon (25) were imported, analyzed, and visualized in HiVis. Only adjacent-normal or control tissues were included in the analysis.

Epithelial-specific genes were defined using a signature matrix obtained from Dan et al. (64).

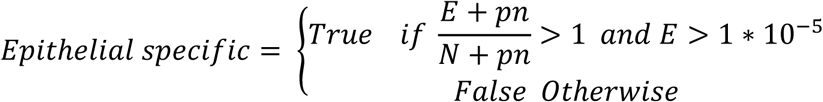

Where *E* is the maximum expression in the enterocytes and goblet cells, *N* is the mean expression across all non-epithelial cell types, and *pn* is the smallest non-zero value in *E*.

Apical, basal and nuclear identities were assigned to bins by manual annotation in QuPath (35). Apical and basal genes were also classified as cytoplasmic. Differential gene expression analysis was performed on epithel-specific genes, excluding mitochondrial genes. Fisher’s exact test was conducted for each sample individually. To integrate results across samples, for each gene log₂FC was taken as the median across all samples, expression level as the mean, and one-sided p-values were combined using Fisher’s method (65). One-sided p-values were concatenated, and corrected for multiple hypothesis testing using the Benjamini-Hochberg method (66), after which they assigned back as one-sided q-values.

For correlation with a previous apical/basal LCM dataset of human small intestine (14), data were imported, and Spearman correlation was calculated on genes with q-value < 0.25.

#### Mouse liver

Raw FASTQ files generated by the sequencer were processed using Space Ranger (v3.1.2, 10x Genomics), including manual image alignment, spatial barcoding, read alignment to the mm10 reference genome, and UMI counting. VisiumHD data from mouse livers were imported, analyzed, and visualized using HiVis. Since the VisiumHD slides contained additional samples from unrelated projects, both the expression data and corresponding images were cropped to the liver regions, and pixels from other samples were excluded. Correlation of expression levels between the two liver samples was assessed using Spearman’s correlation.

##### Single cell segmentation

Single-cell segmentation was performed in QuPath (35) using the Cellpose (37,63) and StarDist (36) extensions. The whole tissue was first segmented using a Random trees pixel classifier, keeping only regions larger than 25 µm^2^, and closing holes smaller than 50 µm^2^. A second pixel classifier was applied to identify the blood vessel regions. Hepatocytes were segmented within the whole tissue mask, using a custom-trained Cellpose model with both ATP1A1 and DAPI channels, and were allowed to expand into blood vessel regions to allow smooth segmentation. The model was trained to delineate the exact borders of hepatocytes, enabling accurate segmentation of mononuclear, binuclear, and cells without nucleus in the imaging plane. Training was performed on 100 regions (120 × 120 µm each) sampled from the dataset. Segmentation performance was evaluated by selecting an additional 20 regions (10 from each sample) and manually segmenting ∼80 cells per region, defined as a ground truth (GT) dataset. Our applied custom Cellpose model was compared against the built-in cyto3 Cellpose model – both based on membrane and nuclei signals – and against nuclei segmentation with expansion approach using Stardist and Cellpose (Supplementary Table 3). To assess segmentation performance an intersection-over-union (IoU) was computed for each pair of matching cells. IoU was computed as the number of intersecting pixels between the two segmented objects divided by the union of the pixels of the two objects. A correct match was considered only if IoU was larger than 0.5, namely if the overlap between the GT and the segmented cell encompassed more than 50% of the pixels in the union of both objects. Following this calculation, commonly used precision metrics were computed, including recall, F1 score Aggregated Jaccard Index (AJI):

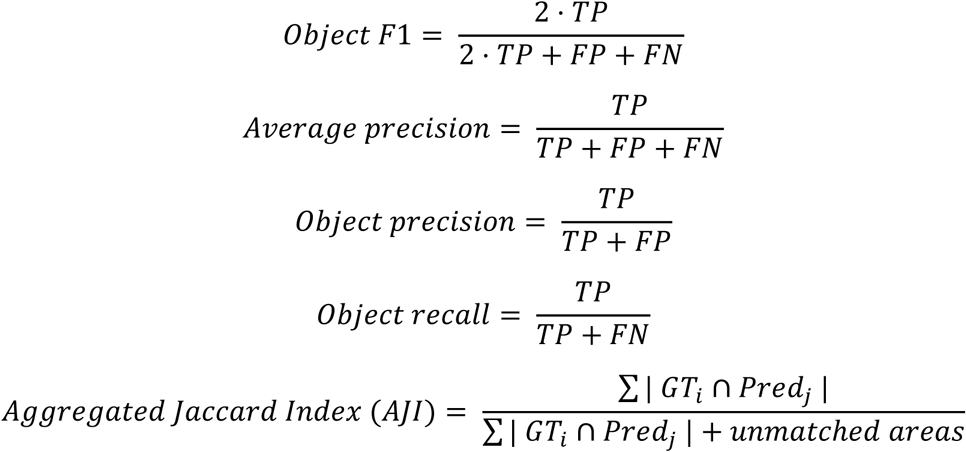

Where TP denotes true positive detections, defined as one-to-one matches between ground truth (GT) objects, and predictions with an IoU ≥ 0.5. FP denotes false positive detections (predicted objects not matched to any GT object), and FN denotes false negatives (GT objects without a corresponding prediction). AJI computes the sum over all matched pairs of their overlapping pixels divided by this sum and the total sum of areas that are unmatched (both non-overlapping pixels in matched pairs and total pixels of unmatched objects and GT).

To allow analyses based on nuclear counts, nuclei were segmented independently using the built-in *cyto3* model, and each nucleus was subsequently matched to its parent cell. NPC<u>s</u> were segmented within blood vessel regions using an in-house–trained StarDist model on the DAPI channel (67), with expansion restricted to vessel regions and capped at 2 µm. The StarDist threshold was set to 0.3 with normalization percentiles of 1 and 99. NPC were further filtered by size (nuclear area 10-80 µm²) and by DAPI intensity >18,000.

##### Lobule zonation analysis

Hepatocyte zonation was inferred using previously published periportal and pericentral marker gene lists (48) by calculating the zonation score 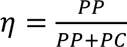, where PP is the sum of the normalized expression of periportal markers and PC is the sum of normalized expression of pericentral markers. The zonation score was binned into six groups using the Fisher-Jenks method (68), and zones were smoothed by calculating the median within a 40 μm radius kernel.

Comparison of gene zonation between samples and with previously published data (49,52) was performed using a center-of-mass (COM) calculation for each gene *g*:

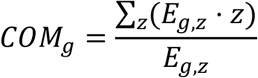

Where *E_g,z_* is the expression level of gene *g* in zone *z*, and *z* is the numeric index of the zone (ranging from 1 to 6). When comparing VisiumHD with the other datasets, the average expression per zone was computed over the pooled cells from both VisiumHD samples prior to COM calculation. The resulting COM values were correlated using Spearman’s correlation.

##### Zonal ploidy probabilities

To determine the probability of hepatocyte ploidy states across zones, nuclear areas were quantified separately for mononucleated and binucleated hepatocytes in each lobular zone. In binucleated cells, the summed areas of both nuclei were considered. For each group, nuclear area distributions were binned into 50 histogram intervals and smoothed (kernel size = 3 bins). Local maxima were detected, and the minimum between the two maxima was used as the decision boundary between diploid and tetraploid nuclei. Mononuclear hepatocytes with nuclear areas below or above this threshold were assigned as 2n or 4n respectively, whereas bi-nucleated hepatocytes with nuclear areas below or above this threshold were assigned as 2n+2n or 4n+4n respectively.

Within each mouse and zone, the proportion of hepatocytes in each ploidy state was calculated as the number of cells in that ploidy class divided by the total number of hepatocytes of the same nuclearity class. These per-mouse proportions were then averaged across mice for each zone and ploidy state.

##### Identification of basal, cortical and nuclear genes

To find large blood vessels and sinusoids between hepatocytes, a pixel classifier was applied with the following parameters:

- Classifier: Random trees
- Resolution: Moderate (2.57 µm/pixel)
- Features:
  - Channels: CD31, ATP1A1, DAPI, autofluorescence
  - Scales: 0.5, 1, 2, 4
  - All available features

The distance from each bin center to the nearest sinusoid pixel was computed. Bin identities were then assigned as follows:

- Blood vessel: Bins with >90% coverage by blood vessel identity
- Basal: bins within 1.5 µm of a blood vessel
- Apical: non-basal bins within 1.5 µm of the cell border
- Cytoplasmic: all remaining non-apical, non-basal bins outside the nucleus

Hepatocyte-specific genes were defined using a signature matrix incorporating all major liver cell types based on Ben-Moshe et al. (69):

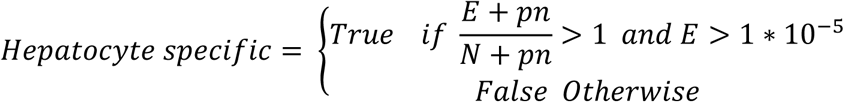

Where *E* is the expression in hepatocytes, *N* is the average expression across all non-hepatocyte cell types, and *pn* the smallest non-zero value in *E*.

Differential gene expression analysis was performed on cells with >500 UMIs, restricted to hepatocyte-specific genes, excluding mitochondrial genes. For basal/cortical analysis, only cells with at least three basal bins and with <20% coverage classified as “blood vessel”, (by the pixel classifier), were included. Fisher’s exact test was applied on each sample individually. To combine both samples, for each gene the log₂FC and expression level were averaged across samples, and one-sided p-values were combined using Fisher’s method (65). One-sided p-values were concatenated, and corrected for multiple hypothesis testing using the Benjamini-Hochberg method (66), after which they assigned back as one-sided q-values.

For correlation with previously published nuclear/cytoplasmic fraction RNA-seq data (18), the data were imported and processed by averaging the two nuclear and two cytoplasmic replicates separately. A log₂(nuclear/cytoplasmic) expression ratio was then computed. Spearman correlation was performed using genes with mean normalized expression >10^-4^ in both datasets and q-value <0.25. To compute the correlation between nuclear to cytoplasmic ratio between central and portal regions, the same test was applied in both zones (central: zones 1,2; portal: zones 5,6), and Spearman’s correlation was calculated on the genes with q-value <0.25 and max normalized expression >10^-5^.

### Human liver

Published data and single-hepatocyte segmentation (44) were imported and analyzed in HiVis. Only one sample (M6), which showed high RNA capture rates, was used in the analysis. Hepatocytes zonation was computed as previously described (44) by calculating the zonation score 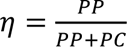, where PP is the summed expression of periportal markers, and PC is the summed expression of pericentral markers. The zonation score was binned into four groups using the Fisher-Jenks method (68), and zones were smoothed by calculating the median within a 50μm radius kernel. Differential gene expression analysis between nuclear and cytoplasmic bins was performed using Fisher’s exact test, restricted to bins in zones 2,3.

## Data availability

The VisiumHD datasets analyzed in this study originate from a combination of publicly available and internal sources. Mouse intestine dataset is available for download from the 10x Genomics website (https://www.10xgenomics.com/datasets). The human liver dataset was obtained from Yakubovsky et al. (44), and the human colon dataset from Oliveira et al., 2025 (25). The mouse liver dataset is available in [ZENODO link will be available upon publication]. <u>Xenium mouse intestine dataset was obtained from</u> <u>Ameku et al.</u> (46)<u>, and available in</u> <u>GSM8695241. Processed Stereo-seq mouse brain dataset was obtained</u> <u>from Qiu et al.</u> (47), <u>raw data is available from Chen et al.</u> (70).

## Code availability

HiVis is available as a Python package via PyPI distribution at https://pypi.org/project/HiVis/. The source code for the HiVis package and the QuPath scripts, together with usage examples, is available in the HiVis GitHub repository: https://github.com/roynov01/HiVis. Detailed documentation is provided at https://hivis.readthedocs.io/latest/. The code used for the analyses in this study is available in [GITHUB link will be available upon publication] and has also been deposited in Zenodo under accession [ZENODO link will be available upon publication].

## Author’s contributions

R.N and S.I conceived the study. R.N and O.G conceived the HiVis tool and performed image analysis. R.N developed the Python code and wrote the documentation. O.G developed the Groovy code. R.N and M.K performed the VisiumHD experiment on mouse liver. T.B and I.G contributed to fluorescence image acquisition. I.K performed the surgeries. I.K, M.F and I.N provided colon samples. S.I supervised the project. R.N and S.I wrote the manuscript. All authors contributed to the manuscript revisions and approved the final version.

## Acknowledgments

We thank Ron Moran for assistance with packaging our tool for distribution via pip. We thank Jacob Elkahal and Eldad Tzahor for providing the CD31 antibody. We thank Elizabeth Vaisbourd for the insightful feedback that improved the clarity and presentation of the figures. S.I. is supported by the Helen and Martin Kimmel Award for Innovative Investigation, the Yad Abraham Research Center for Cancer Diagnostics and Therapy, the Moross Integrated Cancer Center, a Weizmann-Sheba grant, the Israel Science Foundation grants no. 908/21 and 3663/21, the European Research Council (ERC) under the European Union’s Horizon 2020 research and innovation programme grant no. 768956 and a grant from the Ministry of Innovation, Science & Technology, Israel. Optical imaging acquired at the de Picciotto Cancer Cell Observatory In memory of Wolfgang and Ruth Lesser of the Moross Integrated Cancer Center in the department of Life Science Core Facilities, Weizmann Institute of Science.

## Supplementary tables

Supplementary table 1 – Zonation analysis results.

Supplementary table 2 – Subcellular compartments DGE analysis results.

Supplementary table 3 – Segmentation evaluation results.

## Supplementary figures

**Figure S1.**
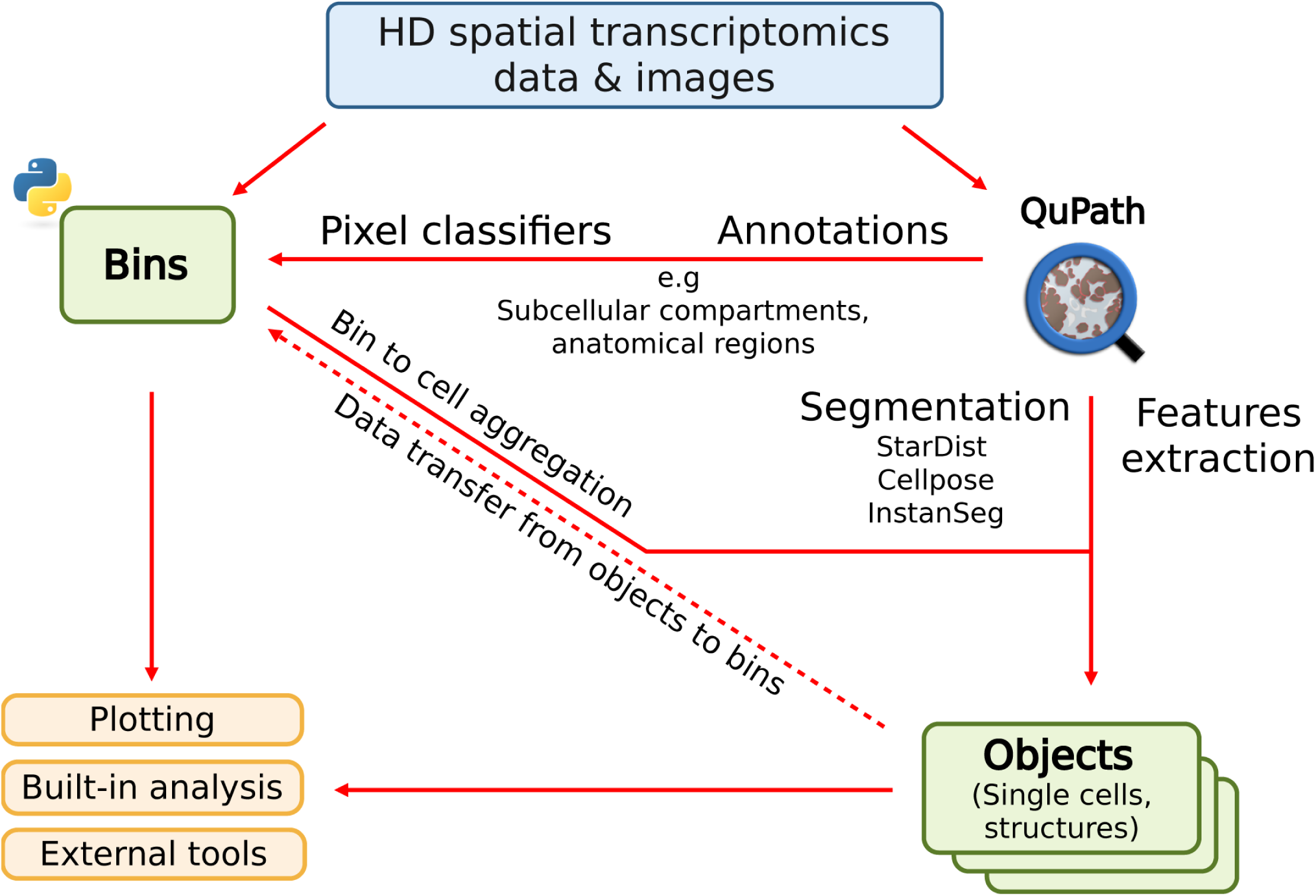
Diagram of the HiVis tool. HiVis is a user-friendly platform for high-definition spatial transcriptomics analysis. Image analysis is performed with QuPath, whereas all other steps are implemented in Python, including aggregation of bins into cells or other objects. HiVis operates at both the bin and aggregation levels, synchronizing data across them. It provides flexible, fully customizable plotting, offers built-in analysis methods, and is compatible with external tools such as Scanpy (39) and Squidpy (40). The diagram was created with BioRender.com.

**Figure S2.**
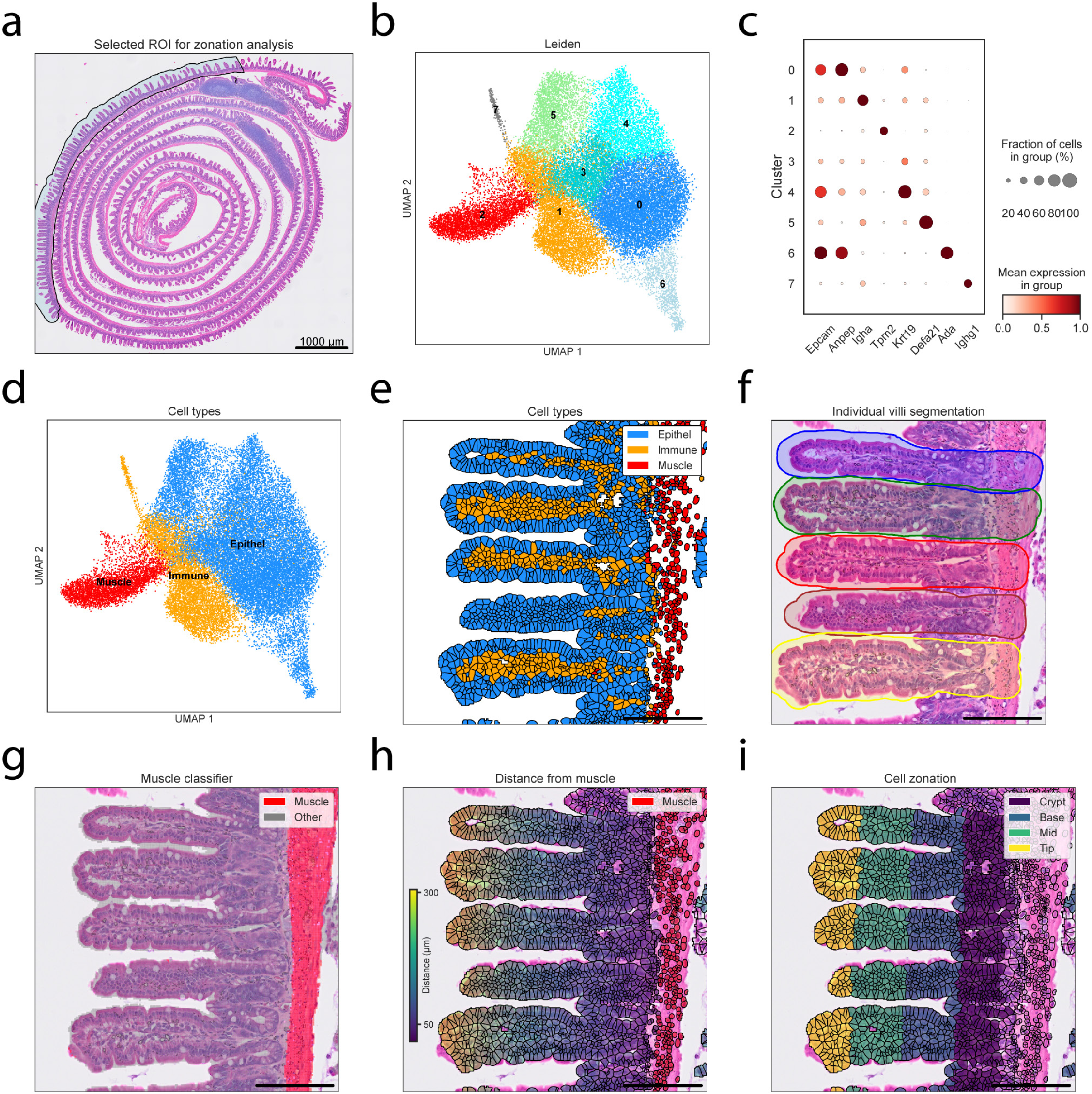
Reconstructing zonation along villi in mouse small intestine. (a) VisiumHD H&E image of mouse small intestine, overlayed with proximal intestine annotation for further analysis. (b) UMAP of single cells colored by leiden clusters. **(c)** Dot plot of normalized gene expression for selected markers across leiden clusters. **(d)** Annotated UMAP of single cells colored by cell type. **(e)** Cells colored by cell type. **(f)** VisiumHD H&E image of mouse proximal intestine, overlayed with individual villi manual annotations. **(g)** Mouse proximal intestine VisiumHD H&E image overlayed with 2×2µm bins colored by muscle identity, based on a pixel classifier. **(h)** Mouse proximal intestine VisiumHD H&E image overlayed with cells colored by the distance from the muscle layer. **(i)** Mouse proximal intestine VisiumHD H&E image overlayed with cells colored by the villus zone. Scale bar in (e-i) – 100µm.

**Figure S3.**
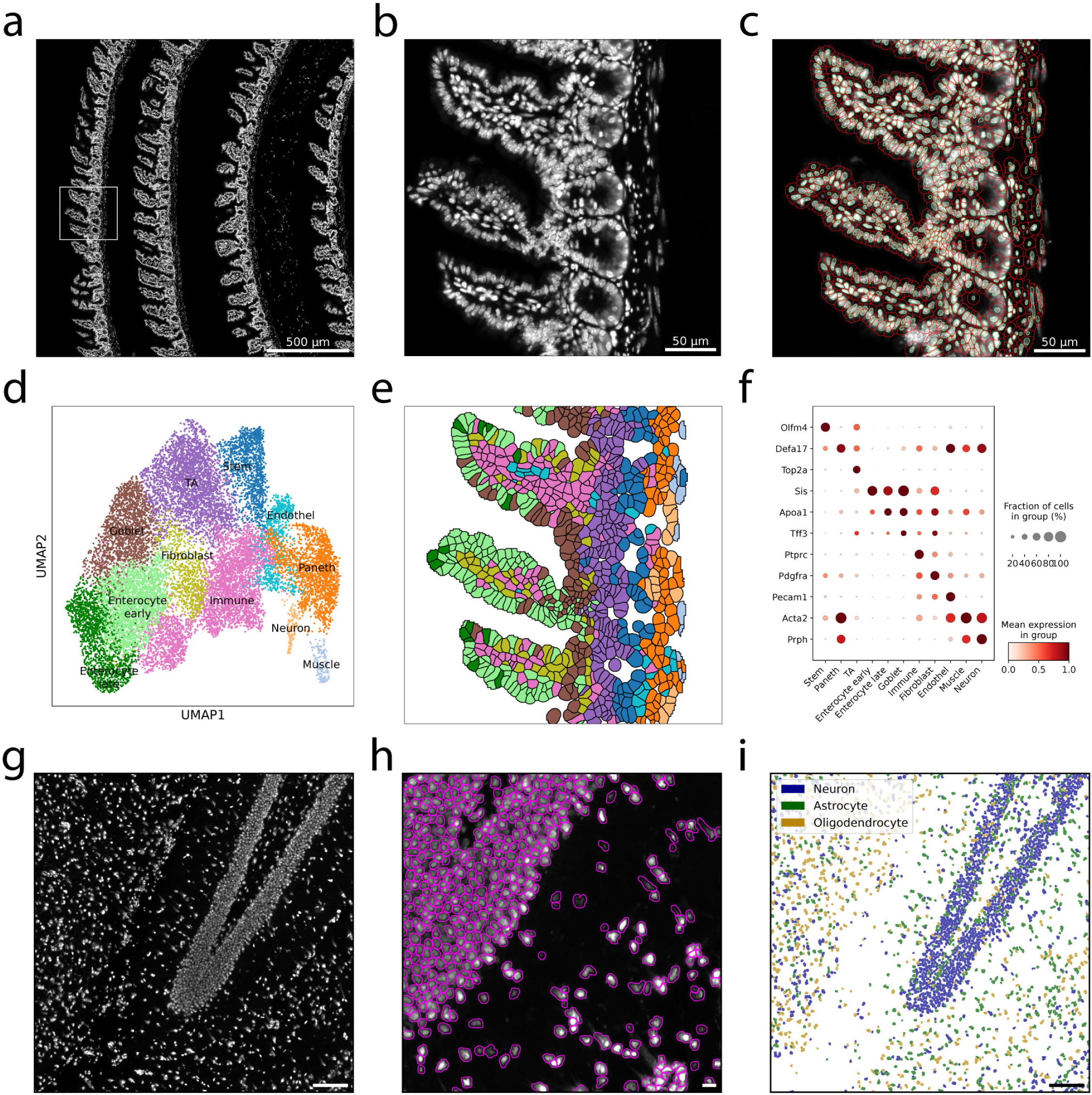
Cross-platform application of HiVis. **(a)** DAPI image of mouse intestine Xenium dataset from Ameku et al. (46). Box denotes selected FOV in (b,c,e). Scalebar 500µm. **(b)** DAPI image of mouse intestine Xenium dataset, selected FOV. Scalebar 50µm. **(c)** Same ROI as (b), with single cells segmented in red. Nuclei in green. Scalebar 50µm. **(d)** Annotated UMAP of single cells colored by cell type. **(e)** Same FOV as (b), with cells colored by cell type. Colors correspond to the cell-type colors in (d). **(f)** Dot plot of normalized gene expression for selected markers across cell types. **(g)** ssDNA image of mouse brain Stereo-seq dataset from Qiu et al. (47). Scalebar 100µm. **(h)** Single cells segmented in mouse brain Stereo-seq dataset. Scalebar 10µm. **(i)** Same FOV as (g) with cells colored by cell-type. Scalebar 100µm.

**Figure S4.**
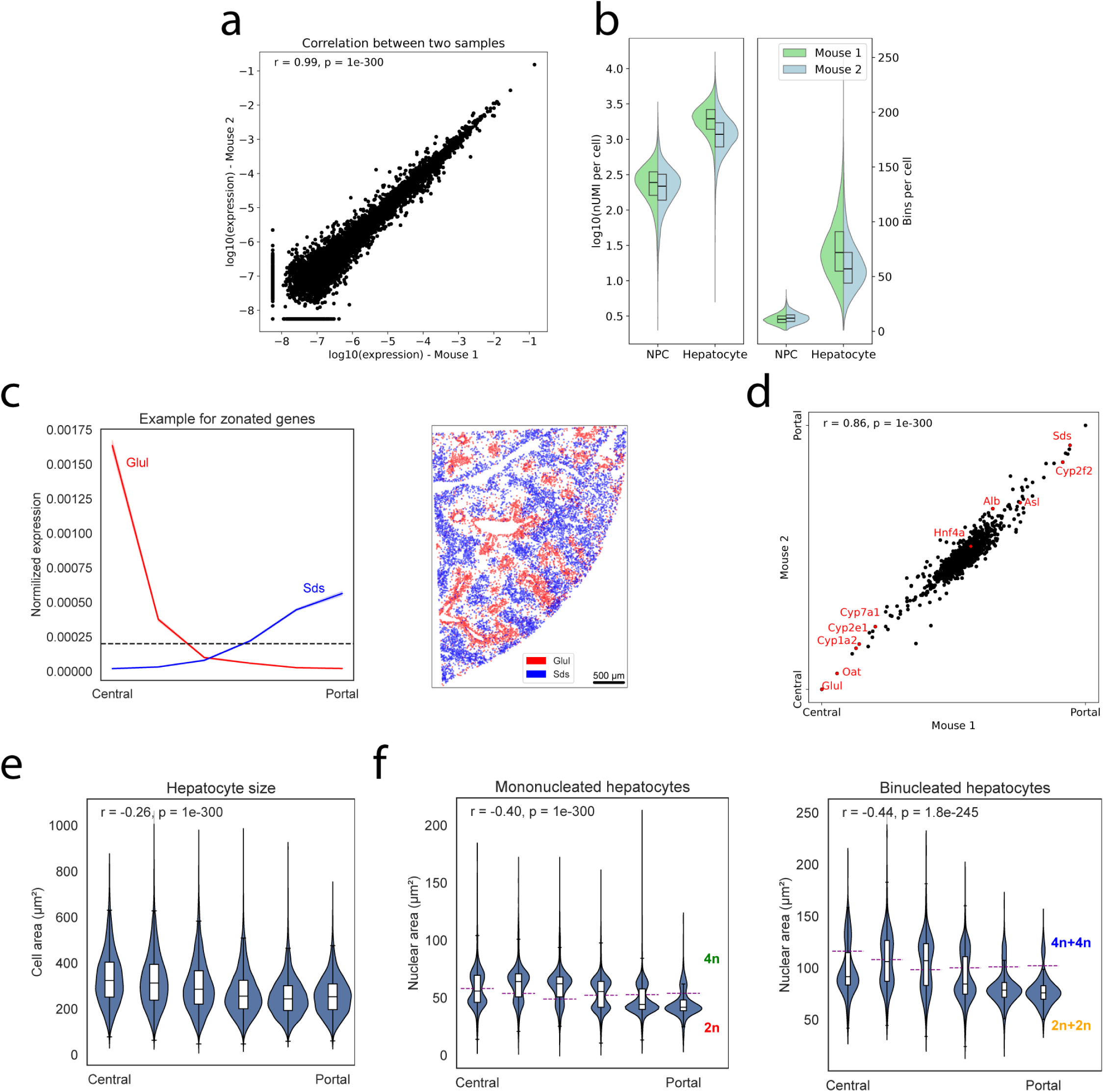
Reconstructing hepatocyte zonation across liver lobules. **(a)** Spearman correlation of normalized gene expression levels between both mice in the experiment. **(b)** Distribution of nUMI (left) and number of 2×2 µm bins (right) in cells from two VisiumHD mouse liver samples. NPC – non parenchymal cells. **(c)** Examples of two zonated genes in single-cell data from mouse liver VisiumHD. Left – normalized expression across liver lobules, data is mean expression, and patch is standard error of the mean. Right – spatial plot of single cells colored by expression of each of the genes. Cells are colored if above the normalized expression > 0.0002. **(d)** Spearman correlation plots of center of mass across liver zones from the two mice in the experiment. **(e)** Violin plots with boxplots showing cell areas of hepatocytes across liver zones. Data is from two samples. Correlation is spearman. **(f)** Violin plots with boxplots showing nuclear area of hepatocytes across liver zones for mononucleated (left) and binucleated (right) hepatocytes. The purple dashed lines mark thresholds separating ploidy classes, indicated by colored labels. Data is from two samples. Correlation is spearman. In boxplots (b, e-f) – horizontal bars are medians, boxes delineate the 25–75 percentiles.

**Figure S5.**
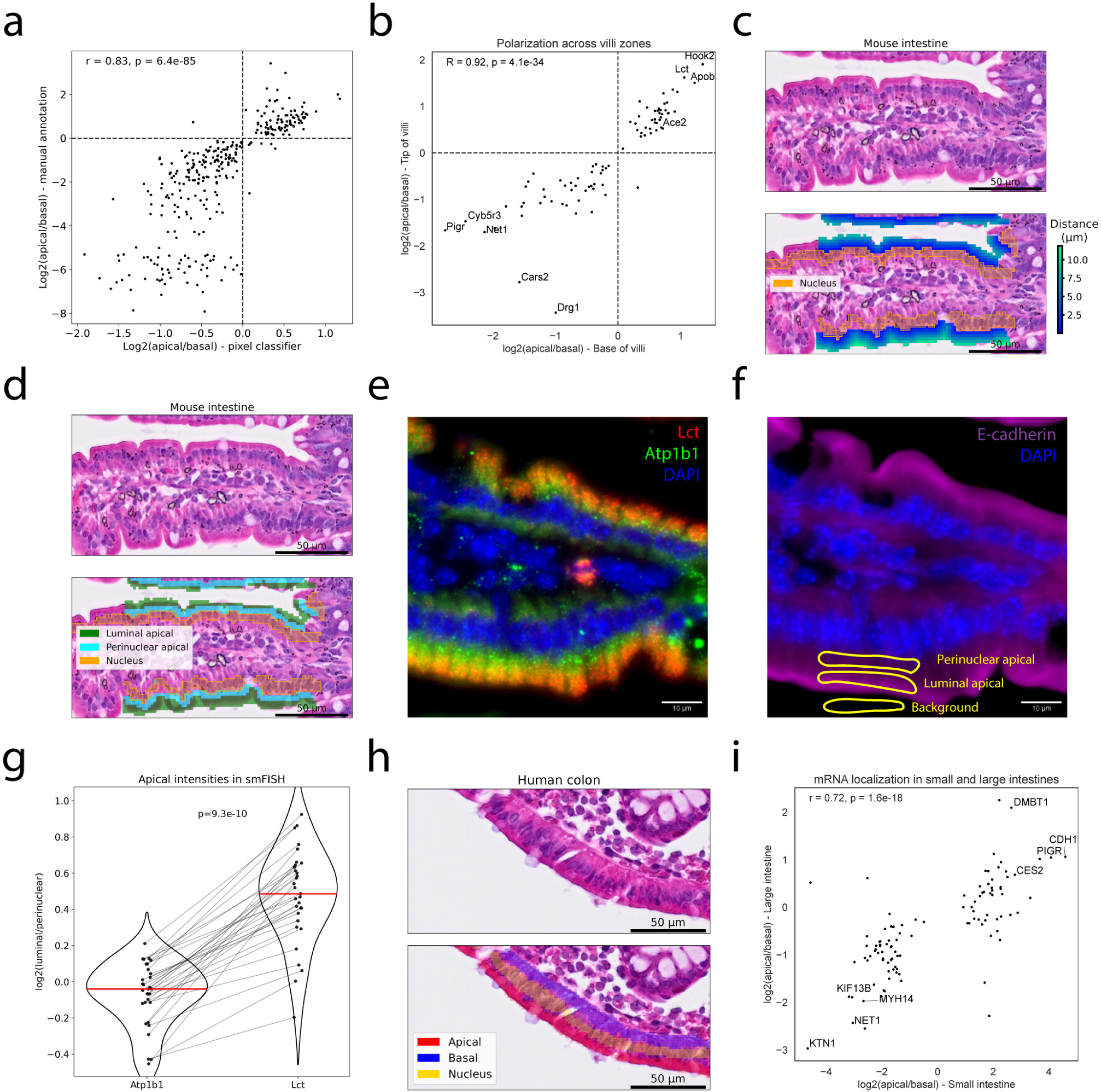
Mapping apical-basal polarization in mouse small intestine and human colon. **(a)** Spearman correlation of apical-basal bias in mouse proximal intestine using manual annotation and pixel classifier. *Genes with normalized expression > 10^-5^ and q<0.05 are used.* **(b)** Spearman correlation of apical-basal bias in base and tip of villi in mouse proximal intestine. Only genes with q<0.05 and normalized expression > 10^-5^ in both zones are used. Selected gene names are shown. **(c)** Mouse proximal intestine VisiumHD H&E image (top) and overlayed with 2×2µm bins colored by the distance to the nuclei (bottom). **(d)** Mouse proximal intestine VisiumHD H&E image (top) and overlayed with 2×2µm bins colored by apical sub-compartments (bottom). **(e)** smFISH staining of Lct and Atp1b1 in mouse proximal intestine. Image is representative of 2 mice. Scale bar, 10 µm. **(f)** Example annotation of apical sub-compartments and background, used for quantification of smFISH intensity. Annotations are based on Immunofluorescence staining of E-cadherin and nuclei. Same FOV as (e). **(g)** Quantification of apical smFISH signal as log2(luminal/perinuclear) intensity for Atp1b1 and Lct. Lines connect paired measurements across annotated apical sub-compartments. Red lines indicate medians. p-value calculated by Wilcoxon signed-rank test. Annotations are from two mice. **(h)** Human colon VisiumHD H&E image (top) and overlayed with 2×2µm bins colored by apical/basal assignment (bottom). **(i)** Spearman correlation of apical-basal bias in colon and Jejunum. Jejunum data is based on previously published LCM RNA-seq (14). Only genes with q<0.25 in both datasets are used. Selected gene names are shown.

**Figure S6.**
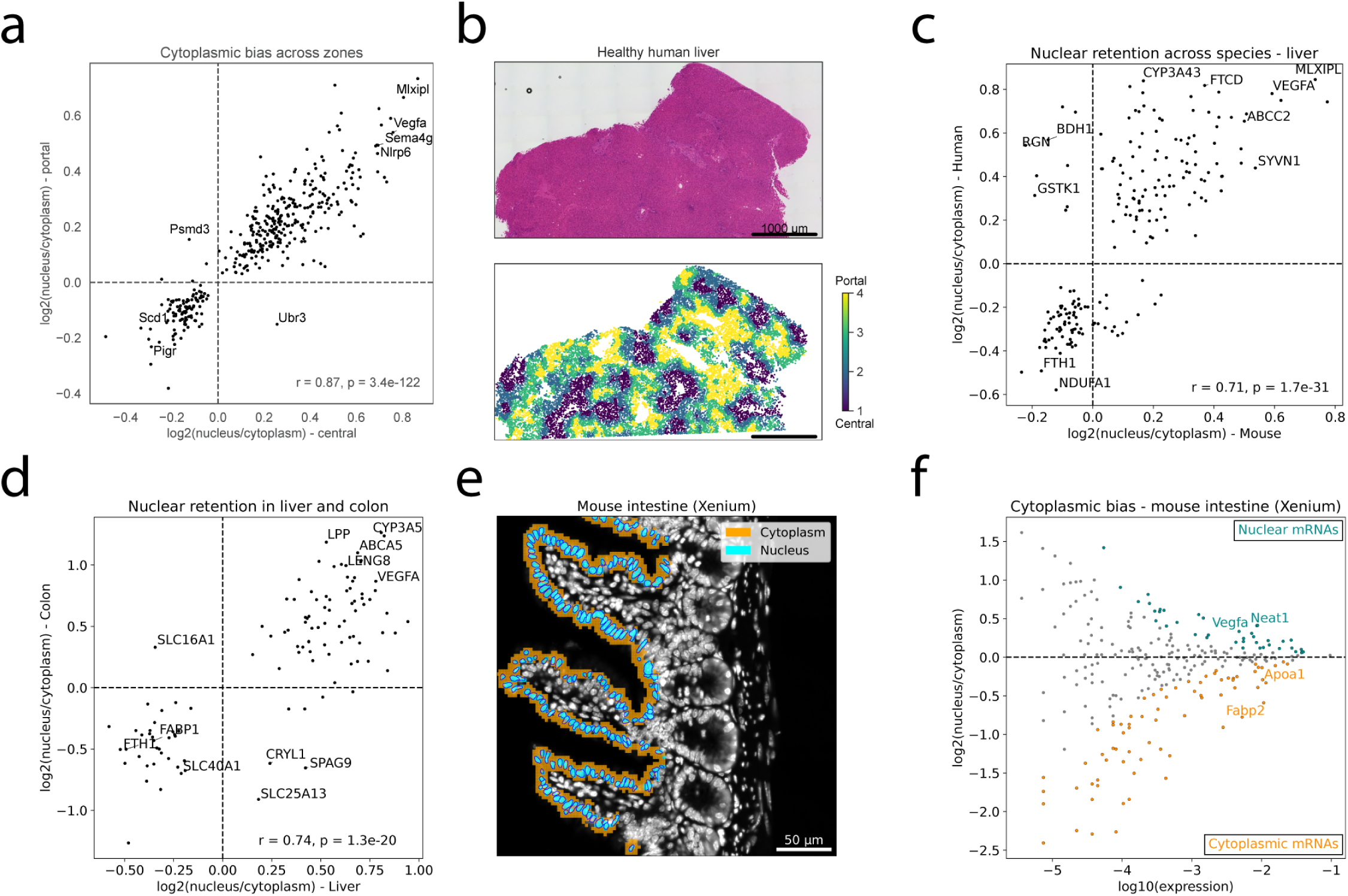
Mapping nuclear retention in livers and Xenium. **(a)** Spearman correlation of nuclear/cytoplasmic bias in pericentral and periportal mouse liver zones. Only genes with q<0.05 and normalized expression > 10^-4^ in both zones are used. Selected gene names are shown. **(b)** Top – human liver VisiumHD H&E image. Bottom – single-cell VisiumHD data of human liver, colored by zones along the central-portal axis. **(c)** Spearman correlation of nuclear/cytoplasmic bias in human and mouse livers. Only genes with q<0.05 and normalized expression > 10^-4^ in both species are used. Selected gene names are shown. **(d)** Spearman correlation of nuclear/cytoplasmic bias in human colon and liver. Only genes with q<0.05 and normalized expression > 10^-4^ in both organs are used. Selected gene names are shown. **(e)** Mouse intestine Xenium (46) DAPI image overlayed with 3×3µm bins colored by nuclear/cytoplasmic assignment, and with nuclear segmentation. **(f)** MA-plot of differential gene expression results in mouse intestine (Xenium). Significant genes are colored cyan and orange (q<0.1). Selected gene names are shown.

